# Canvas: versatile and scalable detection of copy number variants

**DOI:** 10.1101/036194

**Authors:** Eric Roller, Sergii Ivakhno, Steve Lee, Thomas Royce, Stephen Tanner

## Abstract

**Motivation**: Increased throughput and diverse experimental designs of large-scale sequencing studies necessitate versatile, scalable and robust variant calling tools. In particular, identification of copy number changes remains a challenging task due to their complexity, susceptibility to sequencing biases, variation in coverage data and dependence on genome-wide sample properties, such as tumor polyploidy or polyclonality in cancer samples.

**Results**: We have developed a new tool, Canvas, for identification of copy number changes from diverse sequencing experiments including whole-genome matched tumor-normal and single-sample normal re-sequencing, as well as whole-exome matched and unmatched tumor-normal studies. In addition to variant calling, Canvas infers genome-wide parameters such as cancer ploidy, purity and heterogeneity. It provides fast and simple to execute workflows that can scale to thousands of samples and can be easily incorporated into existing variant calling pipelines.

**Availability**: Canvas is distributed under an open source license and can be downloaded from https://github.com/Illumina/canvas.

**Supplementary information**: Supplementary data are available at *Bioinformatics* online.

## INTRODUCTION

The increased throughput of sequencing studies has created high demand for versatile and scalable tools to detect somatic and germline copy number changes. Increasingly complex experimental designs require accurate characterization not only of individual copy number variants (CNVs), but also of global genome and sample properties, such as ploidy, normal contamination and polyclonality (frequently present in cancer samples). Tumors can adopt a wide range of evolutionary strategies depending on the organs involved and treatment protocols, leading to diverse clonal compositions of bulk biopsies (Navin et al., 2010). The interaction of these factors creates an array of different somatic genome architectures that confounds optimization of CNV calling algorithms given an often limited availability of training data. While many CNV calling methods have been introduced, most of them present a number of shortcomings when it comes to scalability and throughput. First, many tools rely on external segmentation software that complicates workflow management and version control. Second, model parameters to infer global genome and sample properties are often hard-coded and difficult to optimize for individual projects. Finally, many tools require the user to provide pre-specified sample-specific parameter values that might not be known prior to CNV detection. For example, TITAN (Ha et al., 2014) and FREEC (Boeva et al., 2011), while inferring heterogeneity and normal contamination respectively, require genome-wide ploidy values as an input, which incorporates a manual step in the variant calling workflow and complicates automation.

We have developed a new tool for CNV calling, Canvas, to address the aforementioned limitations of existing solutions. It fully implements all steps of the variant calling workflow and requires only aligned sequence data and related reference genome files as input. Canvas offers inference of global tumor genome and sample characteristics, including ploidy, purity and heterogeneity, as well as loss of heterozygosity. Alongside fast and easy-to-run whole-genome and exome workflows for both somatic and germline variants, Canvas incorporates a number of global parameter inference strategies based on unsupervised learning approaches such as Expectation Maximization (EM) clustering. This combined functionality makes Canvas a favorable tool for somatic and germline CNV detection in large-scale sequencing projects.

## METHOD

*Outline* The Canvas workflow comprises five distinct modules designed to (1) process aligned read data and calculate coverage bins, (2) perform outlier removal and normalization of coverage estimates, (3) identify segments of uniform copy number, (4) calculate minor allele frequencies (MAF) and (5) assign copy number / allelic states and infer genome-wide parameters. Separate workflows exist for somatic, germline and exome sequencing data. The latter workflow can be run either with or without a matched normal control sample and requires a manifest file with locations of targeted regions. Detailed step-by-step explanations can be found in Supplementary Methods.

*Implementation and performance* Canvas is implemented in C# programming language and can be run on Linux system using mono or on Windows under the .NET framework. A full per-chromosome parallelization is available for all time-consuming modules. An average Canvas runtime for tumor/normal workflow on a 80x/40x coverage matched sample pair and a Linux node with 32 CPUs is 40 minutes (70 minutes when fragment-based GC-content normalization is invoked); with a peak RAM consumption of under 6 gigabytes.

## RESULTS

We tested Canvas on both simulated and real data and also compared its performance with a number of alternative somatic CNV calling tools. The selection of third-party methods was based on the principle that they either showed a superior performance in the previous benchmarks (as in the case of FREEC) or that they offered inference of a number of different genome-wide parameters, like THetA (Oesper et al., 2013). We didn't aim to perform a comprehensive evaluation of CNV callers, as this was covered elsewhere (Alkodsi et al., 2015; Nam et al., 2015). What follows is an overview of test data generation strategies and performance results.

*Simulation data* We have used a haplotype mixing workflow to generate simulation data. Briefly, aligned reads were split into haplotypes inferred from phasing of Platinum Genomes (PG) family (http://www.platinumgenomes.org). These reads along with manually curated CNV calls from previously sequenced genomes at Illumina were used to create truth sets. Simulation also included parameters to generate tumors of different contamination and polyclonality levels.

*Cell lines* Breast carcinoma cell lines HCC2218 and HCC1187 were sequenced to 80x coverage on HiSeq2000 along with matching normal lymphoblastoid cell lines (sequenced to an average of 40X). Exome samples of the same cell lines were prepared using Nextera Rapid Capture Exome reagent kit, and sequenced on a HiSeq 2500. Read titration with normal lymphoblastoid data was used to approximate contamination by normal cells.

*Evaluation strategy and results* A total of 37 samples have been used in benchmarking (Supplementary Results, Section 1). We have focused on exploring concordance between expected and observed copy numbers for each tool. Either exact or directional CN matching was considered. Genome-wide percentage of overlapping regions was used to estimate accuracy, precision and recall. Table 1 shows average metrics values across all simulated and real samples; Supplementary Results provide full details of simulation and evaluation strategies.

**Table 1.**
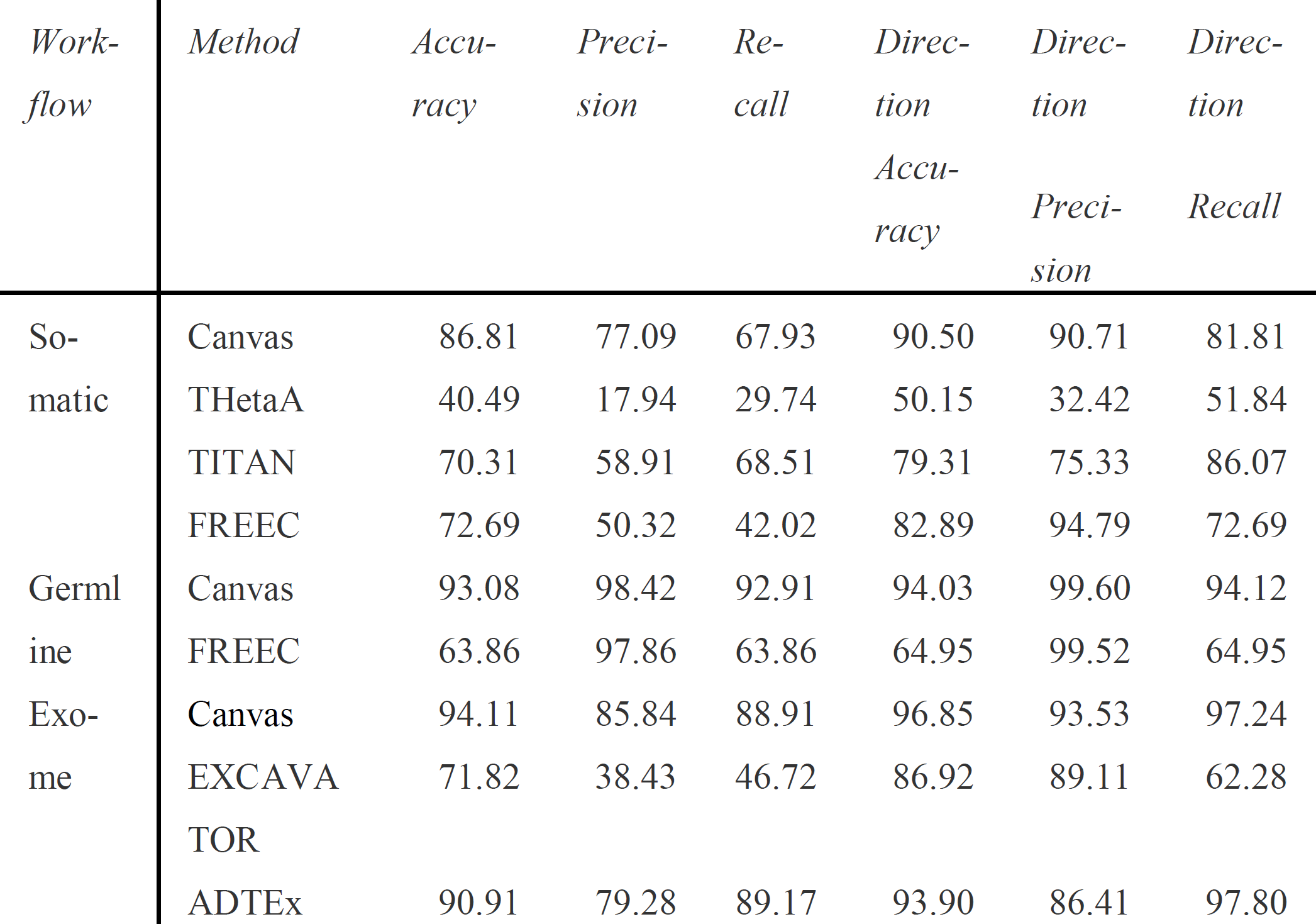
CNV calling performance metrics (average values across all samples for each workflow type)

Canvas attained the largest number of best performing metrics among tools used in comparison for each workflow type (Table 1). For example, in somatic workflow evaluation Canvas had three highest metrics versus two for TITAN and one for FREEC. Moreover, while the distance between Canvas and a second best performing tool was 13.3% for metrics where Canvas performed better, it was only 2.7% for metrics where Canvas didn't show the best result. Moreover, Canvas exhibited a much better performance when exact complimentary between predicted and expected copy number calls was required. Similar observations were made for exome and germline re-sequencing workflows, where Canvas outperformed EXCAVATOR (Magi et al., 2013), ADTEx (Amarasinghe et al., 2014) and FREEC on all but two metrics. Canvas also showed the fastest runtime on a whole genome tumor-normal input in a regular normalization mode and whole genome re-sequencing workflow, completing on average 2.5 times faster than the next fastest method (Supplementary Results, Section 3). To conclude, Canvas is a versatile tool for CNV identification with superior performance metrics across a range of whole-genome and exome sequencing experiments and a fast runtime.

